# *Xanthomonas rydalmerenesis* sp. nov., a novel plant bacteria isolated from *Fragaria x ananassa*

**DOI:** 10.1101/2023.11.12.566340

**Authors:** Daniel J E McKnight, Johanna Wong, Efenaide B Okoh, Fridtjof Snijders, Fiona Lidbetter, John Webster, Mathew Haughton, Aaron E Darling, Steven P Djordjevic, Daniel R Bogema, Toni Chapman

## Abstract

We describe five bacterial isolates that were isolated from *Fragaria x ananassa* in 1976 in Rydalmere, Australia, during routine biosecurity surveillance. Initially, biochemical characterisation identified these isolates as members of the *Xanthomonas* genus. To determine their species, we conducted further analysis using both phenotypic and genotypic approaches. Phenotypic analysis involved using MALDI-TOF MS and BIOLOG GEN III microplates, which confirmed that the isolates belonged to the *Xanthomonas* genus but could not classify species. Genome relatedness indices and extensive phylogenetic analysis confirmed that the isolates belonged to the *Xanthomonas* genus and represented a new species. Based on the absence of virulence factors typically found in *Xanthomonas* spp. genomes, we suggest that these isolates are non-pathogenic. This conclusion was supported by a pathogenicity assay. Based on these findings, we propose the name *Xanthomonas rydalmerenesis*, with DAR34855 = ICMP24941 as the type strain.

## INTRODUCTION

*Xanthomonas* is a genus of Gram-negative proteobacteria that cause disease in over 400 plant species with very few recorded non-pathogens. These hosts include 268 dicotyledonous and 124 monocotyledonous plants, causing significant agricultural impact (1–3). *Xanthomonas* spp. infects a wide variety of economically significant crops, including citrus, cabbage, tomato, capsicum, bean, rice, wheat, and barley (1,2,4). Often, pathogenic bacteria affecting plants consist of a suite of pathovars adapted to infect one host species or closely related species within the same genus, with many exhibiting tissue specificities for the vascular system or mesophyll (5). Symptoms of infections include vascular wilts, blights, cankers, black chaff, black rot, and spots on stems, leaves, and fruit (6). *Xanthomonas* spp. are responsible for significant economic losses worldwide, as they lower the quality of produce and can cause crop loss (7).

*Xanthomonas* spp. use a range of strategies to cause disease in plants. These involve the deployment of virulence factors, including secretion systems, flagella, lipopolysaccharides (LPS), and extracellular polysaccharides (EPS). Secretion system types I to VI have been identified in *Xanthomonas* spp., with the type III secretion system (T3SS) considered the primary virulence determinant. The T3SS is composed of a needle-like macromolecular protein that enables the translocation of exoenzymes, or type 3 (T3) effector proteins, directly into the cytoplasm of the host cell (8). These effector proteins cause cytotoxicity, haemolysis, proteolysis, and phosphorylation/dephosphorylation within the host (9,10). Additionally, *Xanthomonas* spp. use a separate T3SS to secrete extracellular components of the flagella, that it uses for swimming motility. LPS play a critical role in cell wall integrity in Gram-negative bacteria and adherence to plant tissues (11,12). EPS, usually xanthan gum, contributes to the formation of biofilms that enhances adhesion to plant cell surfaces and improves stress tolerance (13).

The identification of species within the *Xanthomonas* genus has a long history. The *Xanthomonas* genus was discovered in 1921 by Ethel Mary Doidge as a pathogen causing canker on tomato and capsicum plants (14). In 1939, Walter John Dowson renamed multiple isolates, previously named *Bacterium vesicatorium* by Doidge, and proposed the genus *Xanthomonas* (15). Isolates were originally classified as belonging to the *Xanthomonas* genus based on phenotypic characteristics. The classification of *Xanthomonas* species was a subject of debate throughout its history due to the limitations of phenetic characterisation methods. Subsequently, these methods were proven to be inaccurate. Many isolates that were believed to belong to the same species were delineated as technological advances were made such as DNA-DNA hybridisation and protein electrophoresis. As a result, the taxonomy of *Xanthomonas* was systematically amended, leading to the reclassification of 20 nomenspecies based on DNA-DNA hybridisation by Vauterin et al. (1995). Genomic sequencing advances have resulted in substantial further reclassifications (17). To designate a distinct novel species, it is necessary to employ diverse methodologies that yield consistent outcomes. Here, we provide carbon utilisation, MALDI-TOF MS, genome relatedness indices, and phylogenetic analysis as evidence to identify and classify a novel *Xanthomonas* species isolated from *Fragaria x ananassa* in Rydalmere, Australia.

## ORIGIN AND ISOLATION

Bacterial samples were collected from *Fragaria x ananassa* growing in Rydalmere, Australia, and submitted to the NSW Department of Agriculture diagnostic laboratories in 1976. They were originally identified as *Xanthomonas* spp. and stored as lyophilised cultures in the NSW Plant Pathology & Mycology Herbarium as specimens DAR34855, DAR34857, DAR34881, DAR34882, and DAR34883. These isolates are referred to as the Rydalmere isolates for the remainder of this article.

## GROWTH AND INITIAL CHARACTERISATION

Bacterial cultures were recovered onto yeast dextrose carbonate (YDC) solid agar and incubated at 25°C for 48 hours. They are Gram-negative, aerobic, and rod shaped. They formed yellow, slightly convex, smooth, circular, mucoid colonies that were 2-3 mm in diameter on YDC media. Isolates were deposited in NZ – ICMP24941

## DNA EXTRACTION, SEQUENCING, AND GENOME ASSEMBLY

DNA extraction for the Rydalmere isolates was performed at the Elizabeth Macarthur Agricultural Institute. Library preparation and Illumina short read sequencing was performed according to Bogema et al. (2018) using a HiSeq2500 system.

To extract DNA for Nanopore sequencing, we retrieved isolates on solid nutrient agar (NA) from glycerol stocks and subsequently sub-cultured onto NA. Single colonies were collected and suspended in 1 mL of PBS. Then samples were centrifuged at 10,000 × g for 1 min and cell pellets were washed 3 × with 1 mL PBS. Genomic DNA was extracted with the Circulomics Nanobind CBB Big DNA Kit (Circulomics, Baltimore, Maryland) according to manufacturer’s instructions. Genomic DNA concentration and extraction quality were determined using a Qubit 4 Fluorometer and Nanodrop 1000 (Thermo Fisher Scientific, Waltham, United States), respectively. Library preparation and Nanopore sequencing was provided by the Garvan Institute of Medical Research, Sydney, Australia. Sequencing libraries were generated using the high molecular weight DNA of each isolate with the Multiplex Ligation Sequencing kit (Nanopore, Cat. Number SQK-MLK111-96-XL). Sequencing was conducted on a PromethION R9.4 flow cell (Nanopore, Cat. Number FLO-PRO002).

Nanopore traces were basecalled with Guppy v6.3.7+532d626 (19) utilising the R9.4.1 super-high accuracy model. Raw reads were filtered to a minimum length of 1000 bp using Filtlong v0.2.1 (20) and draft assemblies were generated using Flye v2.9-b1768, Miniasm v0.3-r179 with Minipolish v0.1.2, Raven v1.8.1, and Necat v0.0.1_update20200803 (21–25). Draft assemblies were combined to produce consensus long-read assemblies using Trycycler v0.5.3 (26). Long-read assemblies were polished using Nanopore reads with Medaka v1.7.1 (27) and with Illumina short reads using two sequential executions of Polypolish v0.5.0 and POLCA from MaSuRCA v4.0.9 (28,29). Whole genome sequences of DAR34855, DAR34857, DAR34881, DAR34882, and DAR34883 were submitted to the NCBI database with the accession numbers CP126170, CP126171, CP126174, CP126173, and CP126172, respectively. Accession numbers for the Illumina short reads and Oxford Nanopore long reads of the Rydalmere isolates are listed in the supplementary table S1.

## GENOME RELATEDNESS INDICES

First, we employed comparative genomic analysis to determine whether the Rydalmere isolates belong to a novel species. Genome relatedness was estimated using average nucleotide identity (ANI) with FastANI v1.32 (30) and digital DNA-DNA hybridisation (dDDH) performed using formula 2 of Genome-to-Genome Distance Calculator (GGDC 3.0) (31).

ANI was calculated to compare the Rydalmere isolates with all 2770 genomes under the *Xanthomonas* genus in the National Center for Biotechnology Information (NCBI) database as of 09/01/2023. Genomes with ANI values greater than the 95% threshold are considered to be the same species (32). ANI values between the Rydalmere isolates were >99.998%, and we concluded that they are clonal. However, < 95% for all but three isolates identified from NCBI: LMG9002 (GCF_009835085.1), 3307 (GCF_014199735.1), and 3498 (GCF_014199795.1). These three NCBI isolates have not previously been classified as a *Xanthomonas* sp. and do not share an ANI >95% with any isolates of known *Xanthomonas* spp. There is no publicly available metadata for 3307 and 3498, but LMG9002 was collected in 1989 from an Orange tree in North America and is non-pathogenic (33).

According to ANI, the most closely related species to the Rydalmere isolates and 3307, 3498, and LMG9002 were *X. sontii, X. sacchari*, and *X. bonasiae* with ANI values of 93.8%, 93.6%, and 88.8%, respectively. These findings are reflected by the dDDH results, which compared DAR34855 to all current *Xanthomonas* type strains that have a publicly available genome. Due to the clonal relationship of the Rydalmere isolates, the results obtained from DAR34855 are representative of the other isolates. All type strains had a dDDH value of <51%, which is lower than the species delineation threshold of 70% (34). The most closely related type strains according to dDDH were *X. sontii, X. sacchari*, and *X. hyacinthi* with values of 50.8%, 48.8%, and 33.3%, respectively. Interestingly, both genomic relatedness indices suggest that *X. fragariae* is the most distantly related *Xanthomonas* sp. to the Rydalmere isolates, despite isolation from the same host (*Fragaria x ananassa*). The results from ANI and dDDH indicate that the Rydalmere isolates described here and three NCBI isolates do not belong to a previously described species. Raw ANI and dDDH values can be found in supplementary tables S2 and S3, respectively.

## PHYLOGENETIC ANALYSIS

The 16S rRNA gene is a widely used molecular marker for the identification and classification of bacterial species. This gene is present in all bacteria and has regions of both conserved and variable sequences that allow for differentiation between bacterial taxa. We extracted the 16S gene sequences from the whole genome sequences using Bakta v1.5.1 and Barrnap v0.9 (35,36). Alignment was performed using MUSCLE v5.1 (37) and a maximum likelihood tree was generated using IQTREE v2.1.4 (38) with 100 bootstrap replicates. The phylogenetic tree was visualised using Interactive Tree of Life (ITOL) (39). All genomes analysed contained a 16S rRNA gene, except *X. arboricola*, which was excluded from the analysis.

Figure 1 demonstrates that the 16S rRNA gene is not a reliable marker for *Xanthomonas* spp., despite its historical use in bacterial species delineation (40). The majority of the *Xanthomonas* type strains are distributed across flat clades, making different species indiscernible. The 16S sequences of the Rydalmere isolates and closely related NCBI isolates are indistinguishable from *X. sontii*. As such, phylogenies generated from other genes or sets of genes is required for bacterial species classification.

**Figure 1.**
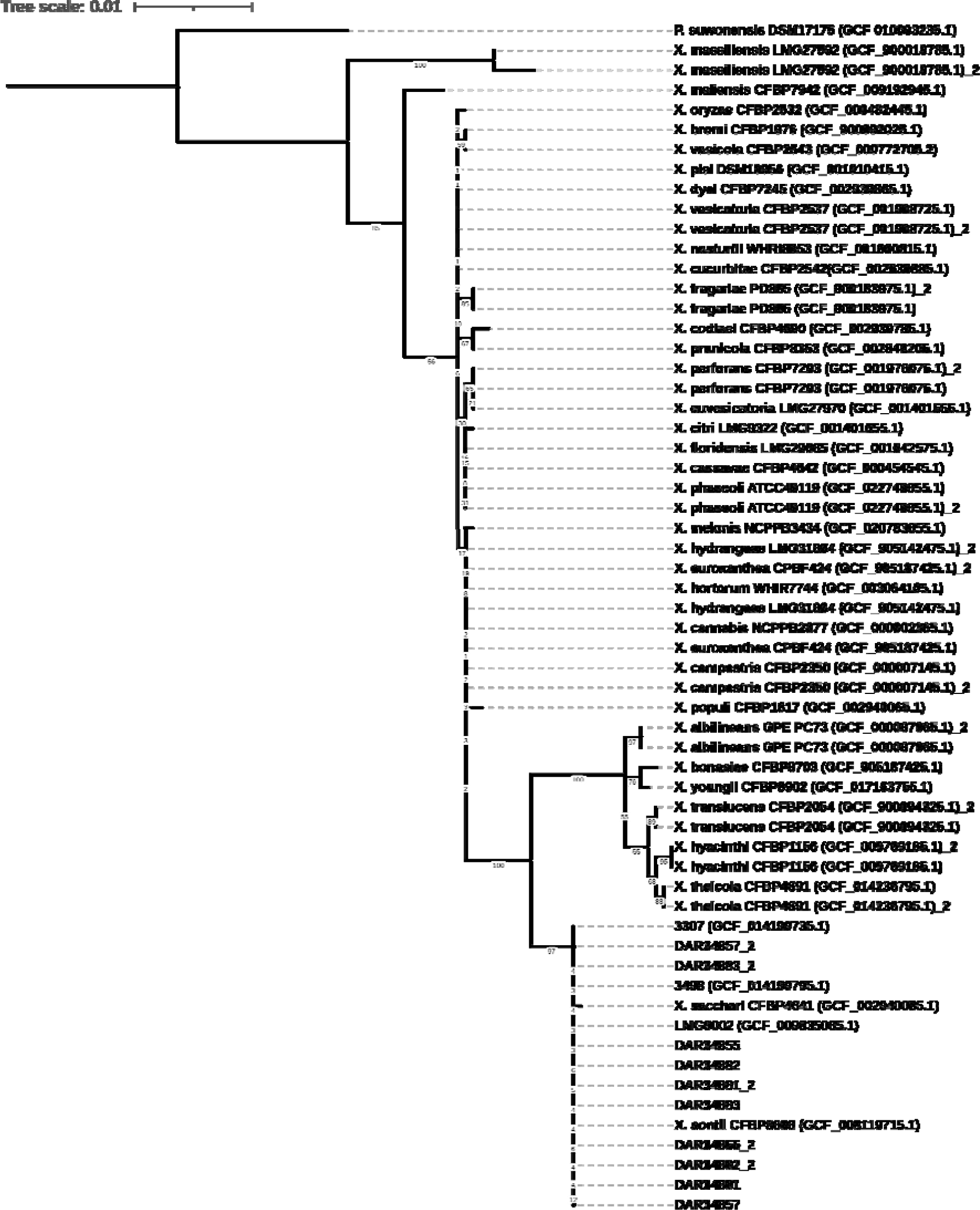
Outgroup rooted maximum likelihood phylogeny of 16S rRNA gene from Rydalmere isolates, three closely related NCBI isolates, all *Xanthomonas* type strains, and *Pseudoxanthomonas suwonensis*. Type strains obtained from NCBI are written as species name followed by type strain and RefSeq database numbers. Genomes with duplicates of 16S rRNA gene are denoted by “_2” for the second copy of the gene.

Gyrase B is a subunit of DNA topoisomerase that plays a crucial role in regulating the supercoiling of DNA during replication and transcription (41). The gene encoding the gyrase B subunit is widely used as a molecular marker for bacterial identification and phylogenetic analysis. This is due to it being highly conserved across bacterial genomes and has sufficient sequence variation to differentiate *Xanthomonas* species (42).

A gyrase B gene tree was generated using sequences from the Rydalmere isolates, three closely related NCBI isolates, all *Xanthomonas* type strains, and *Pseudoxanthomonas suwonensis* as an outgroup (Figure 2). The gene sequences were extracted from the whole genome sequences using Bakta v1.5.1 and Genagr v0.1 (43). All genomes contained a gyrase B gene, excluding *Xanthomonas massiliensis* (GCF_900018785.1). Gene sequences were aligned using MUSCLE v5.1 (37) with default settings and gapped positions were removed using trimaL v1.4.rev22 (44). A maximum likelihood tree with 100 bootstrap replicates was generated using IQTREE v2.1.4 (38) with default settings and was visualised and outgroup rooted using ITOL (39).

**Figure 2.**
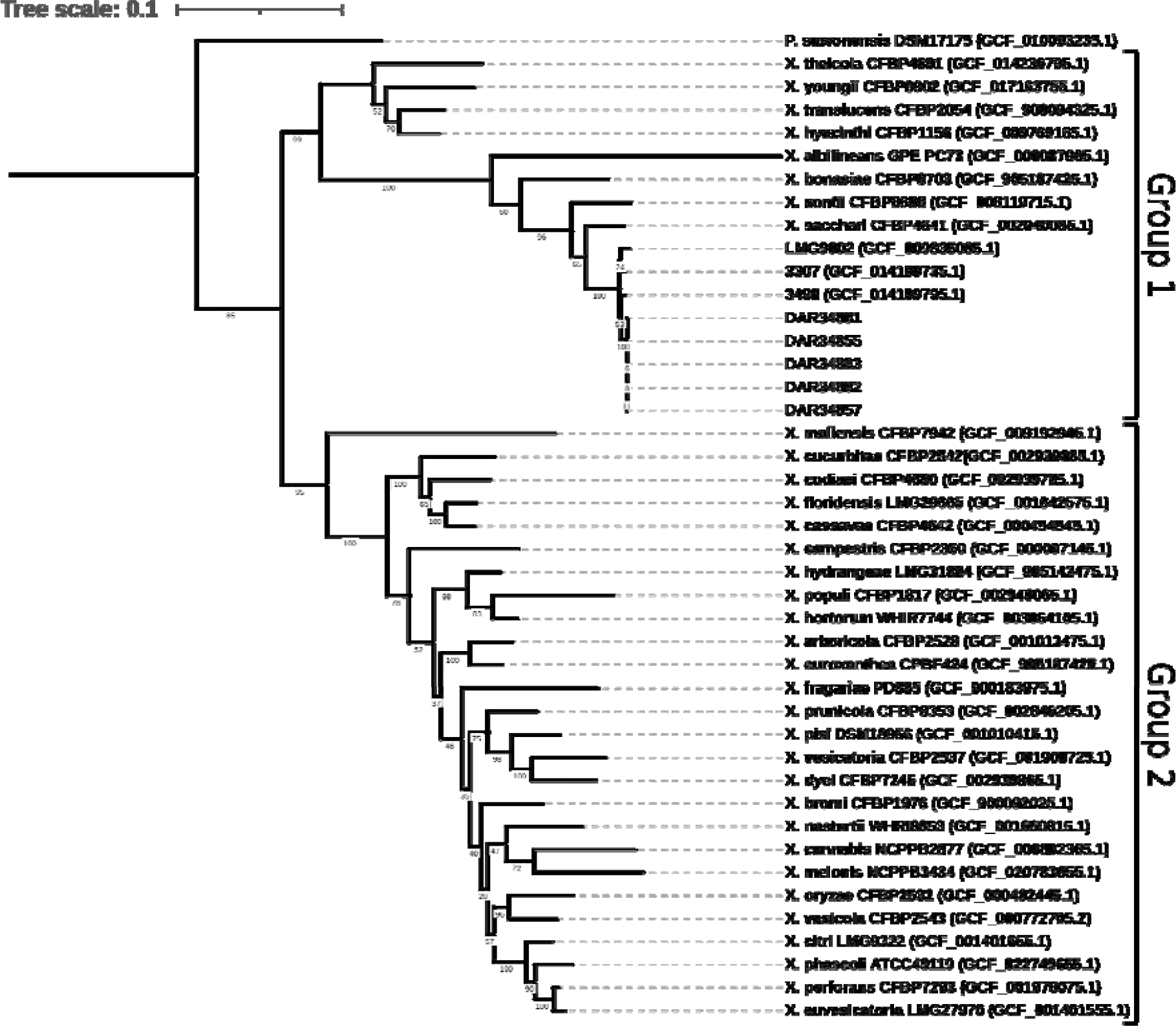
Outgroup rooted maximum likelihood tree using gyrB gene sequences of Rydalmere isolates, three closely related NCBI isolates, all *Xanthomonas* type strains, and *P. suwonensis* as the outgroup. Type strains obtained from NCBI are written as species name followed by type strain and RefSeq database numbers. The two major clades that characterise the *Xanthomonas* genus are labelled according to Young et al. 2008 (45).

The gene tree formed two distinct clades, characteristic of the *Xanthomonas* genus (45–47), with the outgroup external to both groups. Rydalmere and closely related NCBI isolates were positioned within group 1, defined by Young et al. 2008, in a monophyletic sub-clade. *X. sacchari, X. sontii*, and *X. bonasiae* were the nearest species, which reflects the findings of the genome relatedness indices.

Geneious Prime 2023.0.4 (https://www.geneious.com) was used to inspect the sequence alignment. There is a unique 18 nucleotide insertion between 403 and 420 bp in the GyrB alignment that was only detected in the Rydalmere isolates, the three related NCBI isolates, and three closely related *Xanthomonas* species (*X. sacchari, X. bonasiae, X. sontii)*. The insertion does not shift the reading frame.

To further investigate the evolutionary relationship and the accurate phylogenetic position of Rydalmere and closely related NCBI isolates in the *Xanthomonas* genus, a 92-gene multilocus phylogeny was generated using UBCG v3 (48). Analysis included the Rydalmere isolates, three closely related NCBI isolates, all type strains of *Xanthomonas*, and *P. suwonensis* as the outgroup (Figure 3). Concatenated core genes were aligned using MAFFT v7.310 (49) and a maximum likelihood tree was constructed using RaxML (50) with the GTRCAT model. The tree was visualised and outgroup rooted using ITOL (39). The genomes all contained at least 89 of the 92 core genes, with the exceptions of *X. pisi* and *X. phaseoli*, which had 49 and 84, respectively. Accordingly, *X. pisi* was excluded from the analysis.

**Figure 3.**
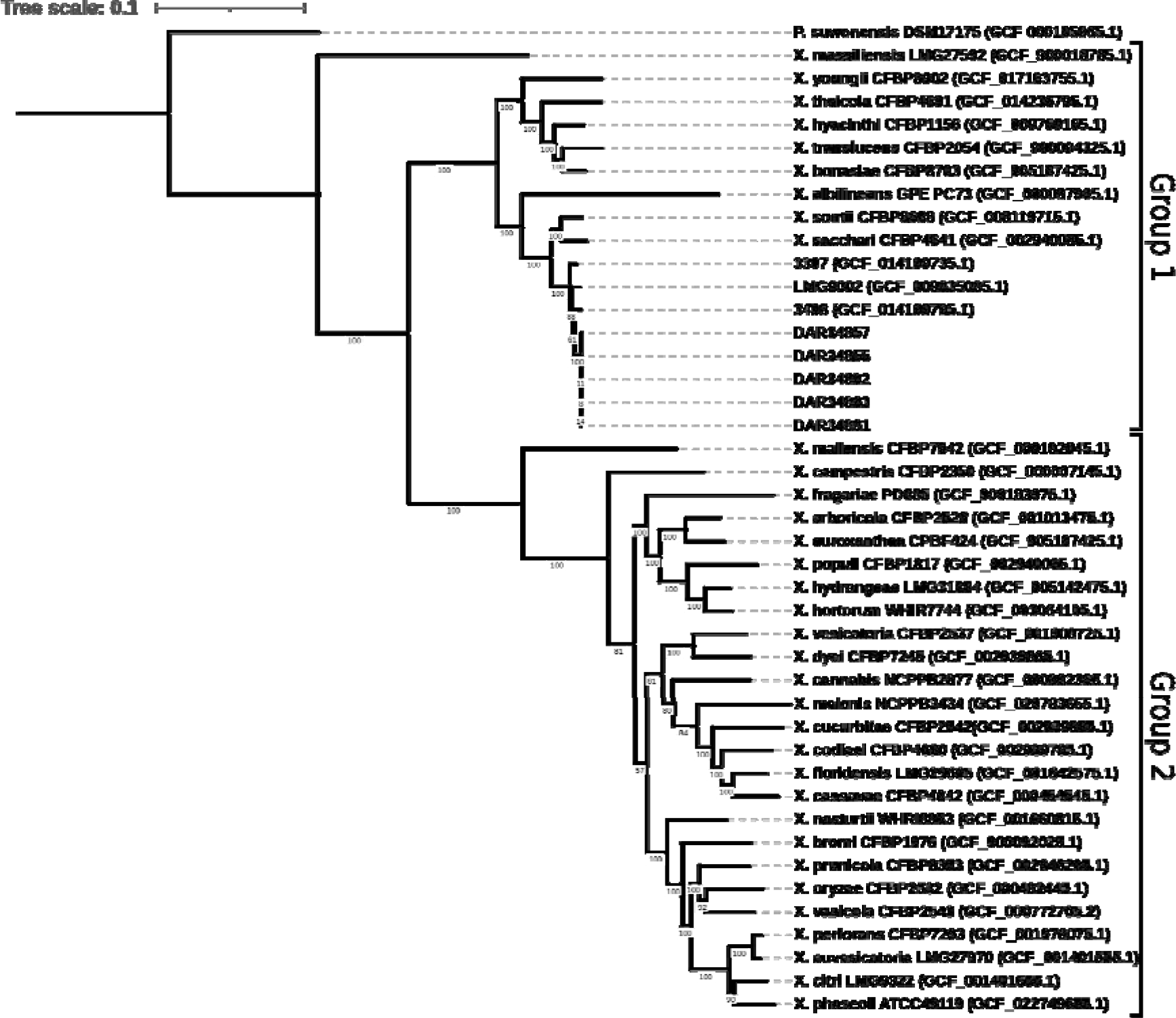
Outgroup rooted maximum likelihood core gene phylogeny of Rydalmere isolates, three closely related NCBI isolates, all *Xanthomonas* type strains, and *P. suwonensis*. Type strains obtained from NCBI are written as species name followed by type strain and RefSeq database numbers. The two major clades that characterise the *Xanthomonas* genus are labelled according to Young et al. 2008 (45).

Analysis revealed that the *Xanthomonas* species grouped into two distinct clades, as previously reported (45,46). The most closely related species was shown to be *X. sacharri* and *X. sontii*, which is consistent with the genomic relatedness indices and *gyrB* gene tree. Rydalmere isolates and similar NCBI isolates formed a monophyletic sub-clade, distinct to that of closely related species. These results reflect the findings from ANI and dDDH analysis that indicate that they do not belong to any known species.

Single nucleotide polymorphisms (SNPs) from the core genomes of the Rydalmere isolates and three closely related NCBI isolates were identified and used to construct an SNP-based phylogeny to investigate their intra-specific relationships (Figure 4). SNPs were detected using Snippy v4.6.0 (51) with default settings and DAR34855 as the reference genome. The alignments were used to generate a maximum likelihood tree with 100 bootstrap replicates using IQTREE v2.1.4 (38). The phylogeny was minimum variance rooted using FastRoot v1.5 (52) and visualised using ITOL (39).

**Figure 4.**
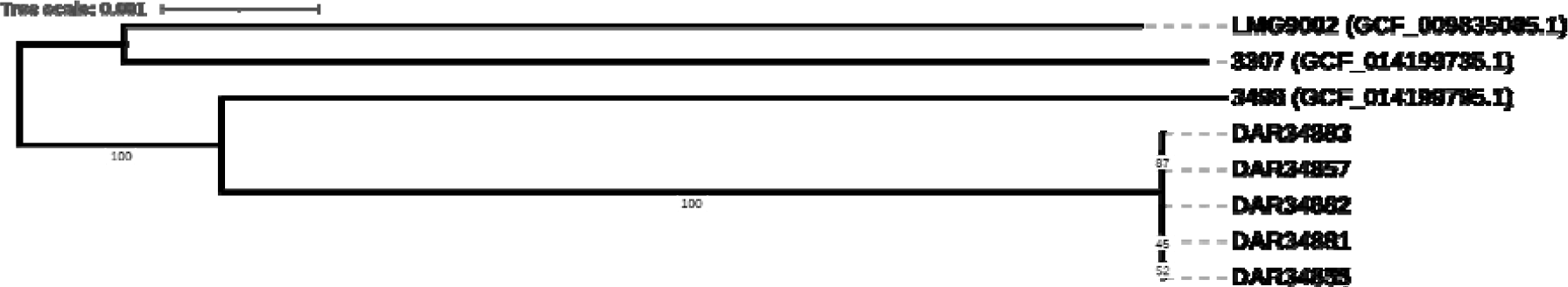
Minimum variance rooted maximum likelihood core genome SNP-based phylogeny of the Rydalmere isolates and three closely related NCBI isolates. NCBI isolates written as strain name followed by RefSeq database number in brackets.

SNPs were only found on five nucleotides between the Rydalmere isolates, which further confirmed their clonality. In comparison, SNPs were detected at 103,591 different nucleotide positions between the Rydalmere isolates and the three closely related NCBI isolates. The Rydalmere isolates exhibit clonality with few SNPs to differentiate them, but in comparison to the NCBI isolates, we found significant genetic variation between members of the species.

## PHENOTYPIC CHARACTERISATION

Phenotypic characterisation of DAR34883 and DAR34855 was performed using Biolog GEN III MicroPlates using protocol A. Selected isolates were sub-cultured from glycerol stock onto solid nutrient-yeast glycerol agar (NYGA) and incubated at 27°C for 48 hours. Then, freshly grown colonies were suspended in inoculating fluid (IF) A using sterile cotton tipped swabs. Density of the inoculating fluid was measured and adjusted to achieve a transmission of 95-98%. A volume of 100 _μ_L of the bacterial suspension was transferred to each well of the MicroPlate and incubated at 30°C. Microplates were read using a MicroStation 2 Reader (Biolog) at 24h, 48h, and 120h. Three plates were used to characterise each of the two isolates with each well measured three times to calculate the mean result. Readings were interpreted as per the manufacturer’s instructions. Carbon utilisation wells were considered positive if they exceeded 160% of the negative control, negative if they were less than 130%, and weak if they fell within the range in between. Chemical sensitivity assay wells were considered positive if the reading was greater than 50% of the positive control. Additionally, if the results between triplicates were inconsistent, they were labelled ‘v’ to signify variability. Values measured at 24h were the most consistent between triplicates and thus was used for analysis.

Across the three replicates, DAR34855 and DAR34883 could utilise 39 substrates, weakly utilise 1, and could not utilise 17 (Table 1; full results supplementary table S4). The remaining substrates were inconclusive as they were variable between isolates. Variation between the isolates was observed as DAR34883 utilised L-Rhamnose as a carbon source, whereas DAR34855 exhibited variable utilisation. Additionally, we found that DAR34883 demonstrated resistance to aztreonam, while DAR34855 was susceptible. Rydalmere isolates differ from closely related species by their ability to utilise L-Galactonic Acid Lactone, 3-Methyl Glucose, and D-Malic Acid. However, they were not unique in their susceptibility/tolerance to different growth environments.

**Table 1.** Phenotypic characterisation of DAR34883 and DAR34885 using BIOLOG GENIII Microplates to determine carbon utilisation, antimicrobial resistance, and growth at varying pH and salt concentrations. Plus (+), minus (-), and ‘v’ signify positive, negative, and variable utilisation of well contents, respectively. *X. hyacinthi, X. sacchari* and *X. theicola* data obtained from BIOLOG GEN III database.

| Characterisation | DAR34855 | DAR34883 | *X. hyacinthi* | *X. sacchari* | *X. theicola* |
| --- | --- | --- | --- | --- | --- |
| Dextrin | + | + | + | + | + |
| D-maltose | + | + | + | + | - |
| D-trehalose | + | + | + | + | + |
| D-cellobiose | + | + | - | + | - |
| Gentiobiose | + | + | - | + | + |
| Sucrose | + | + | + | + | - |
| D-turanose | + | + | - | + | - |
| D-raffinose | - | - | - | + | - |
| α-D-Lactose | + | + | - | + | - |
| D-melibiose | + | + | - | + | - |
| β-Methyl-D- Glucoside | + | + | - | + | - |
| Gelatin | + | + | + | + | + |
| D-fructose | + | + | + | + | + |
| D-galactose | + | + | + | + | + |
| L-fucose | + | + | - | + | + |
| L-rhamnose | v | + | - | - | - |
| D-sorbitol | - | - | - | - | - |
| D-mannitol | - | - | - | - | - |
| D-arabitol | - | - | - | - | - |
| L-alanine | + | v | - | + | - |
| L-aspartic acid | + | + | + | + | + |
| L-glutamic acid | + | + | + | + | + |
| Formic acid | - | - | - | + | - |
| Quinic acid | + | + | - | + | + |
| Citric acid | + | + | + | + | + |
| Bromo-succinic acid | + | v | + | + | + |
| Propionic acid | + | + | - | + | + |
| Acetic acid | + | + | + | + | + |
| L-Galactonic Acid Lactone | + | + | - | - | - |
| 3-Methyl Glucose | + | + | - | - | - |
| D-Malic Acid | + | + | - | - | - |
| L-Rhamnose | v | + | - | - | - |

## PRESENCE AND ABSENCE OF VIRULENCE FACTORS

The presence and absence of *Xanthomonas*-specific virulence factors was performed using BLASTX in Diamond v2.0.15 (53) with no limit on maximum target sequences. A reference database was constructed using 163 previously characterised genes for T3SS, Type III effectors (T3E), flagellar T3SS, Type IV Secretor systems (T4SS), Type VI secretor systems (T6SS), LPS, and EPS from *Xanthomonas* spp. published by Gétaz et al. 2020. Analysis was performed on the Rydalmere isolates and closely related NCBI isolates (Table 2).

**Table 2.**
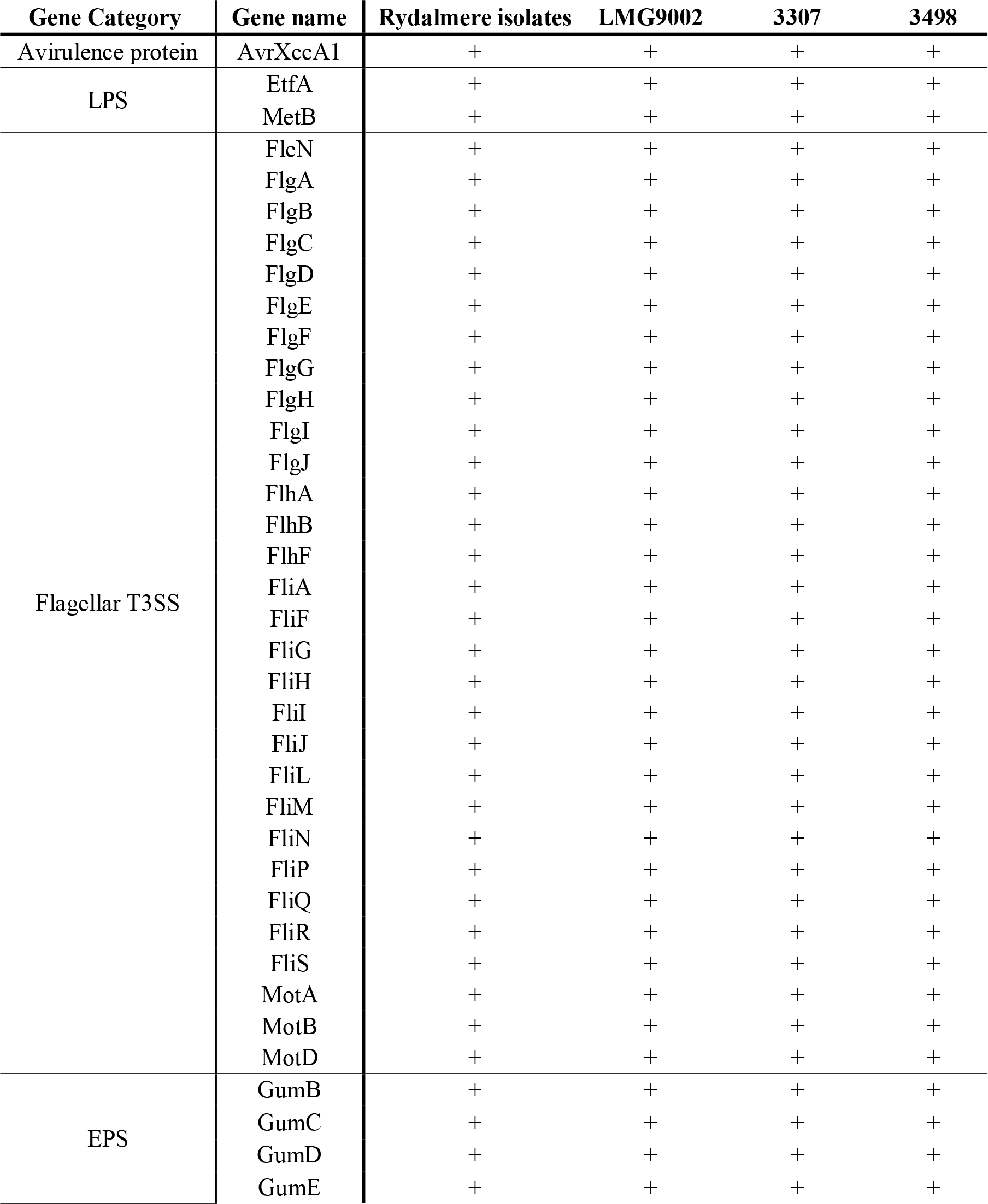
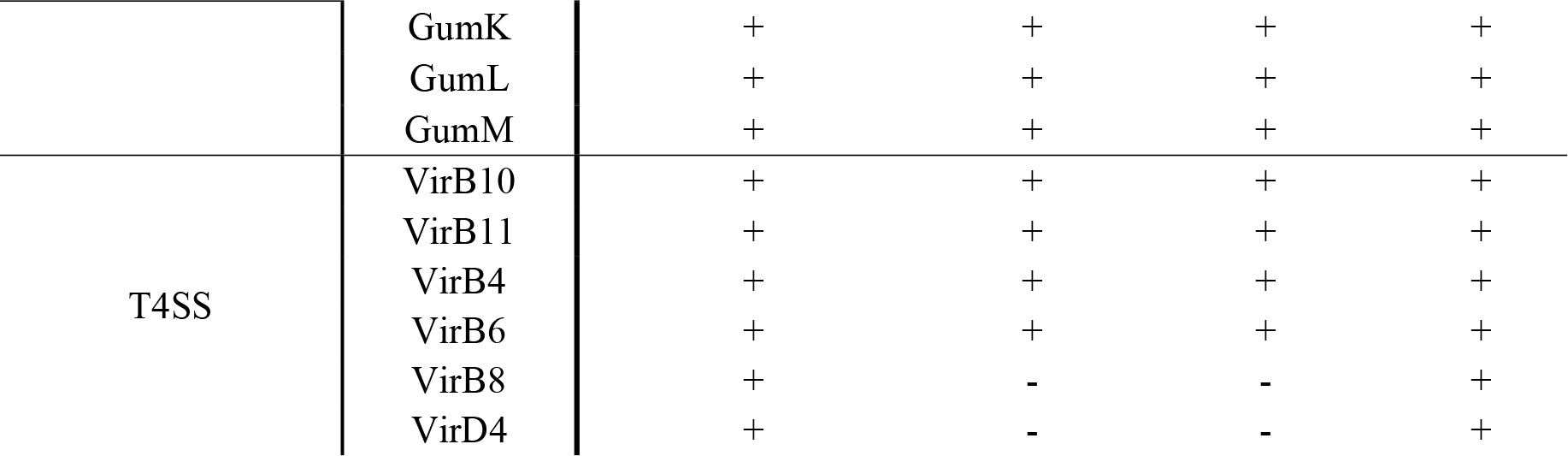
Presence and absence of genes associated with virulence factors in Rydalmere isolates compared to closely related NCBI isolates detected using BLASTX. The presence and absence of a virulence factor is signified by a plus symbol and a minus symbol, respectively. The threshold requirements are a sequence identity similarity of at least 70% and an *e*-value less than 1*e*^-10^.

The Rydalmere isolates, being clonal, had identical virulence factor profiles. Except for the absence of two T4SS genes (VirB8 and VirD4) in LMG9002 and 3307, the virulence gene profiles of the Rydalmere isolates and the three NCBI isolates were nearly identical. We detected the presence of a range of genes that encode the biosynthesis of xanthan, a polysaccharide associated with stress tolerance, host defence inhibition, bacterial adhesion, and biofilm formation (55,56). We detected two genes involved in lipopolysaccharide biosynthesis. All isolates contained a wide variety of flagella T3SS genes, which export the flagella from the cytoplasm to the cell exterior, enabling motility. However, none of the isolates contained the genetic elements required for the synthesis of the virulent T3SS. Additionally, the avirulence factor AvrXccA1 was detected in all isolates. Four to six genes of the T4SS were detected which is involved in the transport of macromolecules associated with virulence (57).

## MALDI-TOF MS ANALYSIS

We generated a profile for DAR34855 using a Bruker Matrix-assisted laser desorption/ionisation-time of flight (MALDI-TOF) Biotyper Microflex LT (Bruker Daltonics, Billerica, MA). The isolate was inoculated onto an NA plate from glycerol stock and incubated for 24-48 hours at 25-28°C. Then, a colony was transferred to an MBT Biotarget plate and an overlay of 1 μL of 70% formic acid was added and allowed to dry. Then, a second overlay of 1 μL of α-Cyano-4-hydroxycinnamic acid was added and allowed to dry before analysis. Finally, the prepared target plate was inserted into the MALDI-TOF Biotyper and was run using settings recommended by Nellessen and Nehl (2023) with a ratio of total-laser-shots to laser-shots-per-raster-spot of 5000:250. Analysis was conducted using the MBT Compass v 5.1.3 software and compared to isolates in our own database and the MBT Compass reference library. The resulting spectrum did not match any known species, with the highest score being another *Xanthomonas* isolate in our database with a log score of 1.83. Log scores between 1.7 and 2.0 are considered ‘low confidence identification’ and thus indicated that this species has not previously been identified and submitted to the Bruker database. The MALDI BioTyper Main Spectra (BTMSP) file for identification of DAR34855 is available in the supplementary file X.rydalmerensisMALDI-TOF.btmsp.zip.

## PATHOGENICITY TESTS

Inoculation of the strawberry cv. elsanta plants with DAR34855 was adapted from the protocol by Kałużna, Kuras & Puławska (2019) in their second trial. Plants were inoculated with 1x10^8^ CFU/mL of the bacteria in sterile water. This concentration was confirmed with CFU plating on NA, incubation, and enumeration. The negative control for each group was inoculated in the same method with sterile water. Each group was a different method of inoculation, totalling three separate methods. Each group (Figure 5) contained four replicate plants and each plant was inoculated three times on each of four different leaves. Negative control plants received comparable treatments, but sterile distilled water was used instead of the bacterial suspension. The pathogenesis trial was run for eight weeks and checked twice weekly for signs of infection, but no hypersensitivity or disease symptoms were observed.

**Figure 5.**
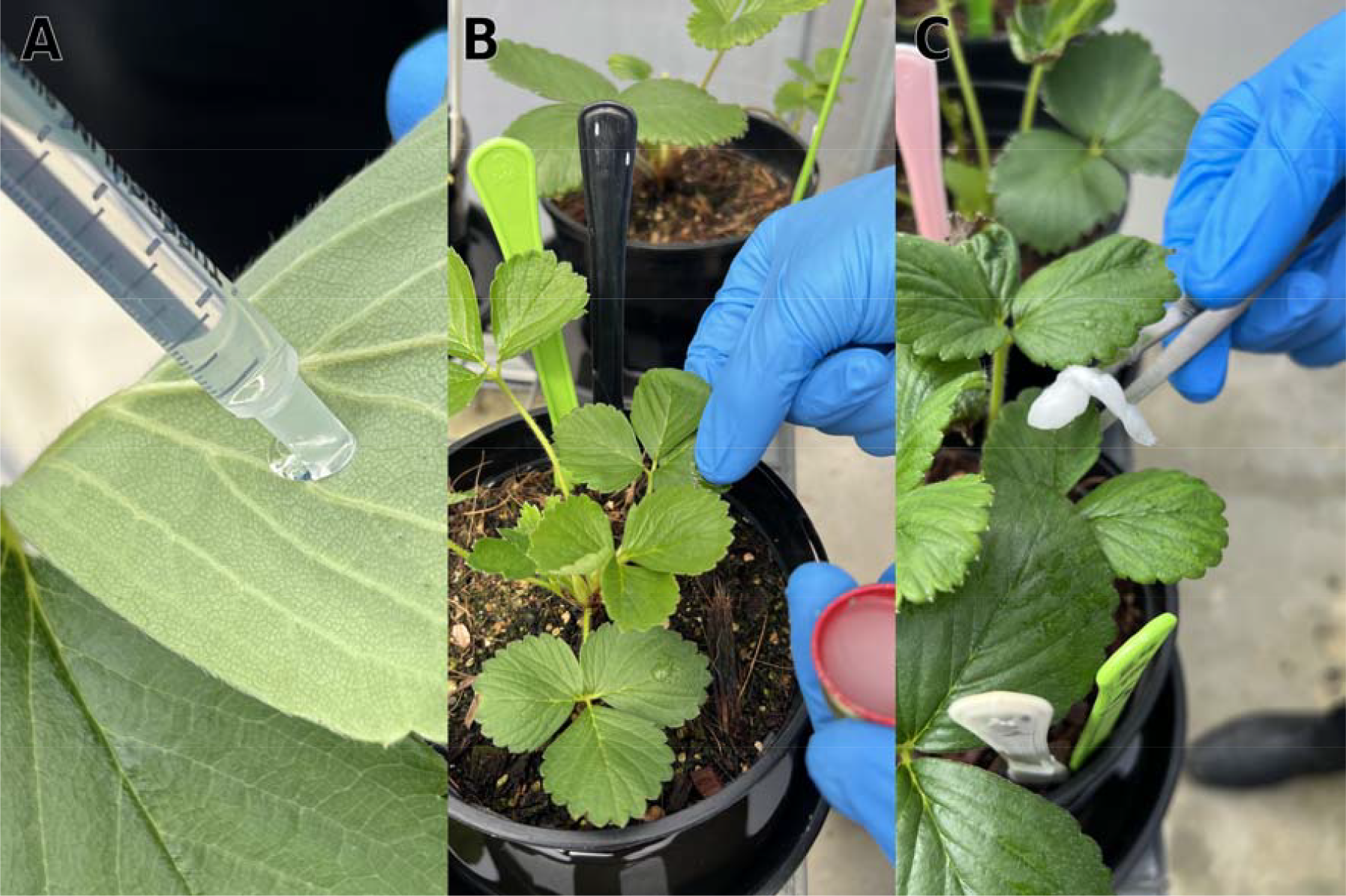
Plant pathogenesis assay inoculation methods. **A** Inoculation of the bacterial suspension using a syringe on the abaxial and adaxial leaf surfaces. **B** Rubbing of leaf surface with a mixture carborundum powder and bacterial suspension. **C** Leaf infiltration using forceps and sterile cotton soaked in the bacterial suspension.

## GENOME FEATURES

Each of the Rydalmere isolates were analysed using QUAST v5.1.0rc1 (59) and Bakta v1.5.1 (35). DAR34855 was designated as the type strain due to its clonal relationship with the remaining four isolates, with the added distinction of being the first collected. The genome of DAR34855 is complete and comprises of a circular chromosome with no plasmids. It is 5.04 Mbp in length, comparable to other species in the genus, with a total number of 4161 coding sequences (CDS). The guanine-cytosine content (GC-content) is 68.7 mol%, which is within the typical range for *Xanthomonas* species.

The core genome of the Rydalmere isolates was defined using Roary v3.13.0 (60) with default parameters, using genome annotations predicted by Bakta v1.5.1 (35). The Rydalmere isolates contain 4165 core genes with 11 shell genes and no cloud genes. When reanalysed with the inclusion of the three closely related NCBI isolates, there were 3524 core genes, 757 shell genes, and 633 cloud genes. No isolates in the analysis were found to contain plasmids.

## DESCRIPTION OF *XANTHOMONAS RYDALMERENSIS* SP. NOV

*Xanthomonas rydalmerensis* (ry.dal.mer.en’sis, O.L. fem. adj. of or belonging to Rydalmere, a suburb of Sydney, where it was first isolated) Cells of the type strain DAR34855 are Gram-negative, aerobic, and rod shaped. It was grown on yeast dextrose carbonate (YDC) solid agar and incubated at 25°C for 48 hours and formed slightly convex, smooth, circular, mucoidal, yellow-pigmented colonies that were 2-3 mm in diameter.

The species is able to utilise α-D-Glucose, Gelatin, Pectin, Tween 40, Dextrin, α-D-Lactose, D-Mannose, D-Galacturonic Acid, D-Maltose, D-Melibiose, D-Fructose, L-Galactonic Acid Lactone, D-Trehalose, β-Methyl-D-Glucoside, D-Galactose, L-Lactic Acid, D-Cellobiose, D-Salicin, 3-Methyl Glucose, Glycerol, L-Aspartic Acid, D-Glucuronic Acid, Citric Acid, Gentiobiose, N-Acetyl-D-Glucosamine, D-Fucose, L-Glutamic Acid, Glucuronamide, α-Keto-Glutaric Acid, Acetoacetic Acid, Sucrose, L-Fucose, D-Fructose-6-PO4, D-Malic Acid, Propionic Acid, D-Turanose, L-Rhamnose, Quinic Acid, L-Malic Acid, and Acetic Acid. It is not pathogenic to strawberry cv. elsanta, its host plant. *X. rydalmerensis*, can be differentiated from other *Xanthomonas* species by dDDH and ANI calculations and with MALDI-TOF spectra. Its genome size is approximately 5.04 Mbp with a DNA GC-content of 68.7 mol%.

The type strain DAR34855 = ICMP24941 was isolated from *Fragaria x ananassa* in Rydalmere, Australia.

## Supporting information

X.rydalmerensisMALDI-TOF.btmsp

## Repositories

Accession numbers for raw reads available in GenBank database.

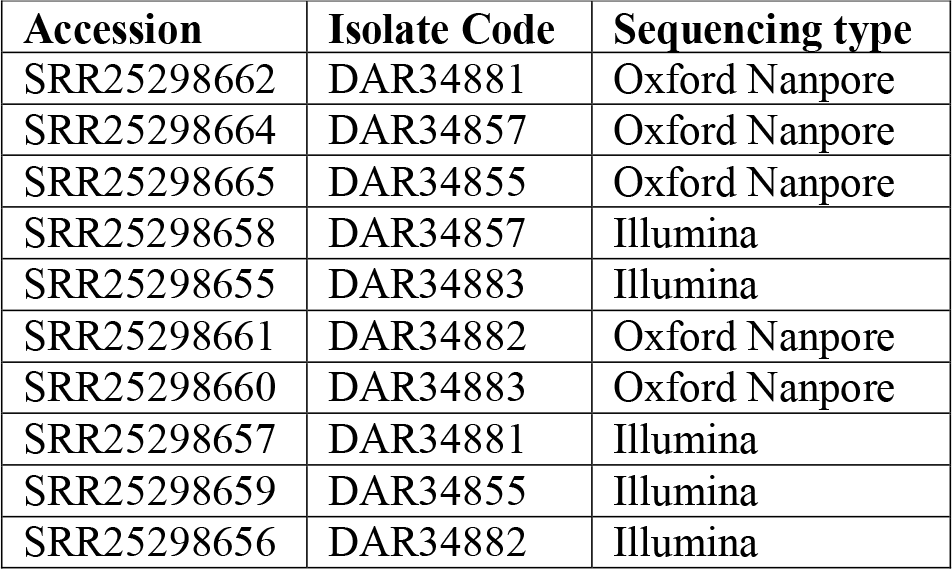

Accession numbers for assembled genomes available in GenBank database

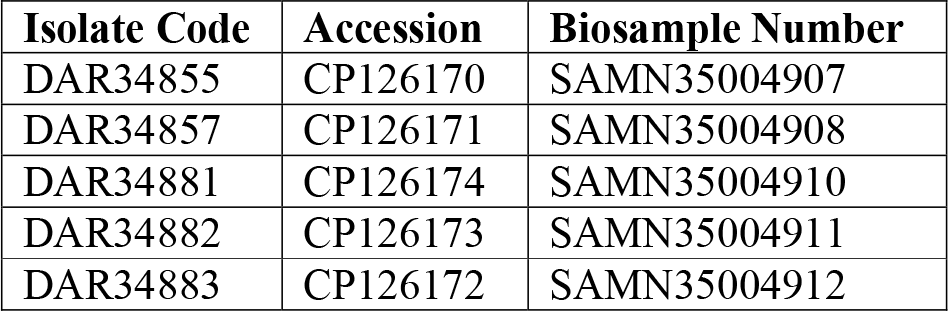

| Isolate Code | Accession | Biosample Number |
| --- | --- | --- |
| DAR34855 | CP126170 | SAMN35004907 |
| DAR34857 | CP126171 | SAMN35004908 |
| DAR34881 | CP126174 | SAMN35004910 |
| DAR34882 | CP126173 | SAMN35004911 |
| DAR34883 | CP126172 | SAMN35004912 |

## FUNDING INFORMATION

This research was funded by the Australian Research Council Linkage project (LP180100593) and Rural Research & Development for Profit/Grains Research and Development Corporation (9177866).

### ACKNOWLEDGEMENTS

We thank Nandita Pathania of the Department of Agriculture, Fisheries and Forestry (Queensland) for help with Biolog data acquisition and the NSW Plant Pathology & Mycology Herbarium for providing cultures. Also, we acknowledge AusGem DPI/UTS for producing high quality Illumina sequencing of the Rydalmere isolates. We acknowledge the financial support provided by the Australian Research Council Linkage project and Rural Research & Development for Profit/Grains Research and Development Corporation. We would like to thank the NSW DPI Advanced Gene Technology Centre for the space and resources for Nanopore sequencing. Finally, we thank Krista Plett and Bernie Dominiak for reviewing this manuscript.

