## Supplementary Data for "*Xanthomonas rydalmerenesis* sp. nov., a novel plant bacteria isolated from *Fragaria x ananassa*"

**Table S1.** NCBI Accession numbers for the reads of the Rydalmere isolates

| **Accession** | **Isolate Code** | **Sequencing type** |
| --- | --- | --- |
| SRR25298662 | DAR34881 | Oxford Nanpore |
| SRR25298664 | DAR34857 | Oxford Nanpore |
| SRR25298665 | DAR34855 | Oxford Nanpore |
| SRR25298658 | DAR34857 | Illumina |
| SRR25298655 | DAR34883 | Illumina |
| SRR25298661 | DAR34882 | Oxford Nanpore |
| SRR25298660 | DAR34883 | Oxford Nanpore |
| SRR25298657 | DAR34881 | Illumina |
| SRR25298659 | DAR34855 | Illumina |
| SRR25298656 | DAR34882 | Illumina |

**Table S2.** Raw ANI values of Rydalmere isolates compared to all *Xanthomonas* type strains and three closely related NCBI isolates.

| **Query** | **DAR34855** | **DAR34857** | **DAR34881** | **DAR34882** | **DAR34883** |
| --- | --- | --- | --- | --- | --- |
| **3307** | 98.2349 | 98.234 | 98.2343 | 98.2351 | 98.2367 |
| **3498** | 98.3681 | 98.3694 | 98.3683 | 98.3731 | 98.3693 |
| **X. albilineans_GPE_PC73_GCF_000087965.1** | 84.2121 | 84.2602 | 84.223 | 84.2563 | 84.2384 |
| **arboricola_CFBP2528_GCF_001013475.1** | 82.8522 | 82.8546 | 82.8532 | 82.8309 | 82.8559 |
| **bonasiae_CFBP8703_GCF_905187425.1** | 88.8607 | 88.8648 | 88.8369 | 88.8659 | 88.8474 |
| **bromi_CFBP1976_GCF_900092025.1** | 81.7237 | 81.698 | 81.7313 | 81.7293 | 81.7208 |
| **campestris_CFBP2350_GCF_000007145.1** | 82.1305 | 82.1237 | 82.129 | 82.1414 | 82.1397 |
| **cannabis_NCPPB2877_GCF_000802365.1** | 82.3088 | 82.2793 | 82.2898 | 82.2943 | 82.3068 |
| **cassavae_CFBP4642_GCF_000454545.1** | 82.3887 | 82.399 | 82.4019 | 82.3932 | 82.4105 |
| **citri_LMG9322_GCF_001401655.1** | 81.953 | 81.9459 | 81.9503 | 81.9672 | 81.9533 |
| **codiaei_CFBP4690_GCF_002939785.1** | 82.6132 | 82.6459 | 82.6046 | 82.645 | 82.5764 |
| **cucurbitae_CFBP2542_GCF_002939885.1** | 82.3965 | 82.3724 | 82.3746 | 82.3582 | 82.3773 |
| **DAR34855** | 100 | 99.9999 | 99.9999 | 99.9999 | 99.9999 |
| **DAR34857** | 99.9998 | 100 | 99.9999 | 99.9999 | 99.9999 |
| **DAR34881** | 99.9999 | 99.9999 | 100 | 100 | 99.9999 |
| **DAR34882** | 99.9999 | 99.9999 | 100 | 100 | 99.9999 |
| **DAR34883** | 99.9999 | 99.9999 | 99.9999 | 99.9999 | 100 |
| **dyei_CFBP7245_GCF_002939865.1** | 81.7075 | 81.7623 | 81.6994 | 81.7321 | 81.706 |
| **euroxanthea_CPBF424_GCF_905187425.1** | 82.9121 | 82.8892 | 82.9162 | 82.9111 | 82.9336 |
| **euvesicatoria_LMG27970_GCF_001401555.1** | 81.9468 | 81.9542 | 81.9389 | 81.9556 | 81.9271 |
| **floridensis_LMG29665_GCF_001642575.1** | 82.13 | 82.1363 | 82.1309 | 82.1562 | 82.1113 |
| **fragariae_PD885_GCF_900183975.1** | 80.922 | 80.9501 | 80.9241 | 80.9107 | 80.9439 |
| **hortorum_WHIR7744_GCF_003064105.1** | 81.9159 | 81.9348 | 81.9096 | 81.9189 | 81.9304 |
| **hyacinthi_CFBP1156_GCF_009769165.1** | 88.6096 | 88.6048 | 88.6259 | 88.6029 | 88.631 |
| **hydrangeae_LMG31884_GCF_905142475.1** | 82.1405 | 82.1574 | 82.1214 | 82.1468 | 82.1359 |
| **LMG9002** | 98.1461 | 98.1469 | 98.146 | 98.1454 | 98.1458 |
| **maliensis_CFBP7942_GCF_009192945.1** | 82.1692 | 82.2029 | 82.2013 | 82.1967 | 82.2029 |
| **massiliensis_LMG27592_GCF_900018785.1** | 83.4443 | 83.4405 | 83.4352 | 83.4269 | 83.4408 |
| **melonis_NCPPB3434_GCF_020783655.1** | 82.4243 | 82.423 | 82.4159 | 82.4417 | 82.4031 |
| **nasturtii_WHRI8853_GCF_001660815.1** | 82.1381 | 82.117 | 82.1401 | 82.1362 | 82.1418 |
| **oryzae_CFBP2532_GCF_000482445.1** | 81.4685 | 81.4621 | 81.459 | 81.4593 | 81.4722 |
| **perforans_CFBP7293_GCF_001976075.1** | 82.0584 | 82.0166 | 82.0375 | 82.036 | 82.0322 |
| **phaseoli_ATCC49119_GCF_022749655.1** | 82.0328 | 82.0109 | 82.0238 | 82.0151 | 82.0328 |
| **pisi_DSM18956_GCF_001010415.1** | 82.1675 | 82.157 | 82.1592 | 82.1655 | 82.1993 |
| **populi_CFBP1817_GCF_002940065.1** | 81.3577 | 81.3749 | 81.3516 | 81.3351 | 81.3501 |
| **prunicola_CFBP8353_GCF_002846205.1** | 81.6171 | 81.598 | 81.621 | 81.6147 | 81.6312 |
| **sacchari_CFBP4641_GCF_002940085.1** | 93.5417 | 93.544 | 93.5412 | 93.5452 | 93.543 |
| **sontii_CFBP8688_GCF_008119715.1** | 93.6715 | 93.6924 | 93.6733 | 93.6964 | 93.6763 |
| **theicola_CFBP4691_GCF_014236795.1** | 87.6236 | 87.593 | 87.6263 | 87.6299 | 87.6066 |
| **translucens_CFBP2054_GCF_900094325.1** | 88.3638 | 88.3604 | 88.3524 | 88.3697 | 88.3522 |
| **vasicola_CFBP2543_GCF_000772705.2** | 81.1839 | 81.2259 | 81.1796 | 81.2246 | 81.2217 |
| **vesicatoria_CFBP2537_GCF_001908725.1** | 81.6552 | 81.6088 | 81.6448 | 81.6293 | 81.637 |
| **youngii_CFBP8902_GCF_017163755.1** | 87.6629 | 87.6532 | 87.6712 | 87.6516 | 87.6796 |

**Table S3.** Raw dDDH values using formula 2 of the X. rydalmerensis type strain against all Xanthomonas type strains

| **Query** | **Reference genome** | **DDH** | **Model C.I.** | **Distance** | **Prob. DDH >= 70%** |
| --- | --- | --- | --- | --- | --- |
| DAR34855 | albilineans_GPE_PC73_GCF_000087965.1 | 28.4 | [26 - 30.9%] | 0.1513 | 0.05 |
| DAR34855 | arboricola_CFBP2528_GCF_001013475.1 | 23.3 | [21 - 25.7%] | 0.1881 | 0 |
| DAR34855 | bonasiae_CFBP8703_GCF_905187425.1 | 32.5 | [30.1 - 35.1%] | 0.1292 | 0.28 |
| DAR34855 | bromi_CFBP1976_GCF_900092025.1 | 22.9 | [20.6 - 25.3%] | 0.1913 | 0 |
| DAR34855 | campestris_CFBP2350_GCF_000007145.1 | 23 | [20.7 - 25.4%] | 0.1907 | 0 |
| DAR34855 | cannabis_NCPPB2877_GCF_000802365.1 | 23.1 | [20.8 - 25.6%] | 0.1891 | 0 |
| DAR34855 | cassavae_CFBP4642_GCF_000454545.1 | 23.2 | [20.9 - 25.6%] | 0.1889 | 0 |
| DAR34855 | citri_LMG9322_GCF_001401655.1 | 23.1 | [20.8 - 25.5%] | 0.1896 | 0 |
| DAR34855 | codiaei_CFBP4690_GCF_002939785.1 | 23.4 | [21.1 - 25.9%] | 0.1868 | 0 |
| DAR34855 | cucurbitae_CFBP2542_GCF_002939885.1 | 22.9 | [20.7 - 25.4%] | 0.1907 | 0 |
| DAR34855 | dyei_CFBP7245_GCF_002939865.1 | 22.7 | [20.4 - 25.1%] | 0.193 | 0 |
| DAR34855 | euroxanthea_CPBF424_GCF_905187425.1 | 23.4 | [21.1 - 25.9%] | 0.1868 | 0 |
| DAR34855 | euvesicatoria_LMG27970_GCF_001401555.1 | 23.7 | [21.4 - 26.1%] | 0.1846 | 0 |
| DAR34855 | floridensis_LMG29665_GCF_001642575.1 | 23.2 | [20.9 - 25.7%] | 0.1883 | 0 |
| DAR34855 | fragariae_PD885_GCF_900183975.1 | 22.5 | [20.2 - 24.9%] | 0.195 | 0 |
| DAR34855 | hortorum_WHIR7744_GCF_003064105.1 | 22.9 | [20.6 - 25.4%] | 0.1911 | 0 |
| DAR34855 | hyacinthi_CFBP1156_GCF_009769165.1 | 33.3 | [30.8 - 35.8%] | 0.1258 | 0.37 |
| DAR34855 | hydrangeae_LMG31884_GCF_905142475.1 | 23 | [20.7 - 25.4%] | 0.1905 | 0 |
| DAR34855 | maliensis_CFBP7942_GCF_009192945.1 | 23 | [20.7 - 25.4%] | 0.1905 | 0 |
| DAR34855 | massiliensis_LMG27592_GCF_900018785.1 | 23.3 | [21 - 25.7%] | 0.1878 | 0 |
| DAR34855 | melonis_NCPPB3434_GCF_020783655.1 | 23.2 | [20.9 - 25.6%] | 0.1887 | 0 |
| DAR34855 | nasturtii_WHRI8853_GCF_001660815.1 | 23.1 | [20.8 - 25.5%] | 0.1899 | 0 |
| DAR34855 | oryzae_CFBP2532_GCF_000482445.1 | 22.7 | [20.4 - 25.1%] | 0.193 | 0 |
| DAR34855 | perforans_CFBP7293_GCF_001976075.1 | 23.1 | [20.8 - 25.5%] | 0.1898 | 0 |
| DAR34855 | phaseoli_ATCC49119_GCF_022749655.1 | 22.9 | [20.7 - 25.4%] | 0.1908 | 0 |
| DAR34855 | pisi_DSM18956_GCF_001010415.1 | 23 | [20.7 - 25.5%] | 0.1901 | 0 |
| DAR34855 | populi_CFBP1817_GCF_002940065.1 | 22.5 | [20.2 - 25%] | 0.1946 | 0 |
| DAR34855 | prunicola_CFBP8353_GCF_002846205.1 | 22.9 | [20.6 - 25.4%] | 0.1911 | 0 |
| DAR34855 | sacchari_CFBP4641_GCF_002940085.1 | 48.8 | [46.2 - 51.4%] | 0.0745 | 15.89 |
| DAR34855 | sontii_CFBP8688_GCF_008119715.1 | 50.8 | [48.2 - 53.5%] | 0.0697 | 21.45 |
| DAR34855 | theicola_CFBP4691_GCF_014236795.1 | 32.2 | [29.8 - 34.7%] | 0.1307 | 0.25 |
| DAR34855 | translucens_CFBP2054_GCF_900094325.1 | 32.1 | [29.7 - 34.6%] | 0.1313 | 0.24 |
| DAR34855 | vasicola_CFBP2543_GCF_000772705.2 | 22.5 | [20.2 - 24.9%] | 0.1949 | 0 |
| DAR34855 | vesicatoria_CFBP2537_GCF_001908725.1 | 22.6 | [20.3 - 25%] | 0.1943 | 0 |
| DAR34855 | youngii_CFBP8902_GCF_017163755.1 | 30.5 | [28.1 - 33%] | 0.1395 | 0.13 |

**Table S4.** All phenotypic Biolog GEN III microplate data for DAR34883 and DAR34855 with closely related Xanthomonas species. X. indica (PPL560) data retrieved from Rana et al. 2022. X. sacchari and X. theicola data obtained from BIOLOG GEN III database

| **Characterisation** | **DAR34883** | **DAR34855** | **X. indica** | **X. sacchari** | **X. theicola** |
| --- | --- | --- | --- | --- | --- |
| **Negative Control** | - | - | - | - | - |
| **D-Raffinose** | - | - | - | + | - |
| **α-D-Glucose** | + | + | + | + | + |
| **D-Sorbitol** | - | - | - | - | - |
| **Gelatin** | + | + | + | + | + |
| **Pectin** | + | + | + | + | - |
| **p-Hydroxy- Phenylacetic Acid** | - | - | - | - | - |
| **Tween 40** | + | + | + | + | + |
| **Dextrin** | + | + | + | + | + |
| **α-D-Lactose** | + | + | + | + | - |
| **D-Mannose** | + | + | + | + | + |
| **D-Mannitol** | - | - | - | - | - |
| **Glycyl-L-Proline** | v | v | v | + | + |
| **D-Galacturonic Acid** | + | + | v | + | - |
| **Methyl Pyruvate** | v | v | + | + | + |
| **γ-Amino-Butryric Acid** | v | - | - | - | + |
| **D-Maltose** | + | + | + | + | - |
| **D-Melibiose** | + | + | + | + | - |
| **D-Fructose** | + | + | + | + | + |
| **D-Arabitol** | - | - | - | - | - |
| **L-Alanine** | v | + | + | + | - |
| **L-Galactonic Acid Lactone** | + | + | - | - | - |
| **D-Lactic Acid Methyl Ester** | - | - | - | - | - |
| **α-Hydroxy- Butyric Acid** | v | - | v | + | - |
| **D-Trehalose** | + | + | + | + | + |
| **β-Methyl-D- Glucoside** | + | + | + | + | - |
| **D-Galactose** | + | + | + | + | + |
| **myo-Inositol** | - | - | - | - | - |
| **L-Arginine** | v | - | - | - | - |
| **D-Gluconic Acid** | - | - | - | - | - |
| **L-Lactic Acid** | + | + | + | + | + |
| **β-Hydroxy-D,L- Butyric Acid** | v | v | v | - | + |
| **D-Cellobiose** | + | + | + | + | - |
| **D-Salicin** | + | + | + | + | - |
| **3-Methyl Glucose** | + | + | - | - | - |
| **Glycerol** | + | + | v | + | + |
| **L-Aspartic Acid** | + | + | + | + | + |
| **D-Glucuronic Acid** | + | + | v | + | - |
| **Citric Acid** | + | + | + | + | + |
| **α-Keto-Butyric Acid** | v | w | v | + | + |
| **Gentiobiose** | + | + | + | + | + |
| **N-Acetyl-D- Glucosamine** | + | + | + | + | + |
| **D-Fucose** | + | + | - | + | - |
| **D-Glucose- 6-PO4** | v | v | v | - | - |
| **L-Glutamic Acid** | + | + | + | + | + |
| **Glucuronamide** | + | + | v | + | + |
| **α-Keto-Glutaric Acid** | + | + | v | + | + |
| **Acetoacetic Acid** | + | + | v | + | + |
| **Sucrose** | + | + | + | + | - |
| **N-Acetyl-β-D- Mannosamine** | v | v | v | - | - |
| **L-Fucose** | + | + | v | + | + |
| **D-Fructose- 6-PO4** | + | + | v | + | - |
| **L-Histidine** | v | w | v | - | - |
| **Mucic Acid** | - | - | - | - | - |
| **D-Malic Acid** | + | + | - | - | - |
| **Propionic Acid** | + | + | + | + | + |
| **D-Turanose** | + | + | + | + | - |
| **N-Acetyl-D- Galactosamine** | - | - | - | + | - |
| **L-Rhamnose** | + | v | - | - | - |
| **D-Aspartic Acid** | - | - | - | - | - |
| **L-Pyroglutamic Acid** | - | - | - | - | - |
| **Quinic Acid** | + | + | + | + | + |
| **L-Malic Acid** | + | + | + | + | + |
| **Acetic Acid** | + | + | + | + | + |
| **Stachyose** | - | - | - | - | - |
| **N-Acetyl Neuraminic Acid** | - | - | - | - | - |
| **Inosine** | - | - | - | - | - |
| **D-Serine** | w | w | - | + | - |
| **L-Serine** | v | + | + | + | - |
| **D-Saccharic Acid** | - | - | - | - | - |
| **Bromo-Succinic Acid** | v | + | + | + | + |
| **Formic Acid** | - | - | - | + | - |
| **Positive Control** | + | + | + | + | + |
| **1% NaCl** | + | + | + | + | + |
| **1% Sodium Lactate** | + | + | + | + | + |
| **Troleandomycin** | - | - | v | - | - |
| **Lincomycin** | + | + | + | + | + |
| **Vancomycin** | + | + | v | + | - |
| **Nalidixic Acid** | - | - | - | - | + |
| **Aztreonam** | - | + | v | + | - |
| **pH 6** | + | + | + | + | + |
| **4% NaCl** | - | - | v | + | - |
| **Fusidic Acid** | + | + | v | - | + |
| **Rifamycin SV** | + | + | + | + | + |
| **Guanidine HCl** | - | - | - | - | - |
| **Tetrazolium Violet** | + | + | + | + | - |
| **Lithium Chloride** | - | - | - | + | - |
| **Sodium Butyrate** | - | - | - | - | - |
| **pH 5** | - | - | v | + | - |
| **8% NaCl** | - | - | - | - | - |
| **D-Serine** | - | - | - | - | - |
| **Minocycline** | - | - | - | - | - |
| **Niaproof 4** | v | + | v | + | - |
| **Tetrazolium Blue** | + | + | + | + | - |
| **Potassium Tellurite** | - | - | v | - | - |
| **Sodium Bromate** | - | - | - | - | - |
